# Estimating narrow-sense heritability from genome-wide data in admixed populations

**DOI:** 10.1101/581389

**Authors:** Georgios Athanasiadis, Doug Speed, Mette K. Andersen, Emil V. R. Appel, Niels Grarup, Ivan Brandslund, Marit Eika Jørgensen, Christina Viskum Lytken Larsen, Peter Bjerregaard, Torben Hansen, Anders Albrechtsen

**Author notes:** Correspondence (G.A.), (A.A.).

## Abstract

Finding an efficient framework for estimating total narrow-sense heritability in admixed populations remains an open question. In this work, we used extensive simulations to evaluate existing linear mixed model frameworks in estimating total narrow-sense heritability in two population-based cohorts from Greenland and compared the results to data from unadmixed individuals from Denmark. When our analysis focused on Greenlandic sib pairs, the model with two relationship matrices, one capturing identity by descent and one capturing identity by state, returned heritability estimates close to the true simulated value, while using each of the two matrices alone led to downward biases. When phenotypes correlated with ancestry, heritability estimates were inflated. Based on these observations, we propose a post-estimation PCA-based adjustment that recovers successfully the true simulated heritability. We use this knowledge to estimate the heritability of ten quantitative traits from the two Greenlandic cohorts and report differences such as lower heritability for height in Greenlanders compared to Europeans. In conclusion, narrow-sense heritability in admixed populations is best estimated using a mixture of genetic relationship matrices on individuals with at least one first-degree relative included in the sample.

## Introduction

Heritability is the fraction of phenotypic variance attributed to genetics. More specifically, assuming that the variance 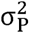 of a phenotype equals the sum of its genetic 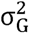 and environmental variance 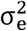, heritability in its broad sense (*H*^*2*^) is expressed by the ratio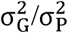.^1,2^ The genetic variance 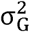 can be further broken down into its additive 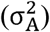, dominant 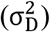, and epistatic 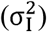 components.^3,4^ Most of the existing methods focus on the fraction of phenotypic variance 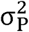 owing to additive effects alone 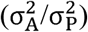 – the so-called narrow-sense heritability (*h*^2^).

Heritability is interesting in its own right, but it is also pivotal in quantitative genetic studies with many practical uses. Because, by definition, heritability measures the contribution of genetics to a phenotype, it allows us to gain insights into the genetic architecture of a trait. Moreover, knowledge about the heritability of a trait helps us evaluate the effectiveness of a genome-wide association study (GWAS), as the so-called single nucleotide polymorphism (SNP) heritability 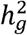 of a trait informs us about the maximum discovery potential of a given genotyping platform. Similarly, heritability estimates provide an upper bound to the accuracy of polygenic predictions with predictors having higher performance potential in traits with higher heritability.

There exist many ways to estimate narrow-sense heritability *h*^*2*^ and they usually boil down to estimating the genetic variance owing to additive effects 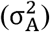. Assuming an additive model, a classical approach is to use the phenotypic correlation between related individuals,^1,2,5^ which for a pair j and k is

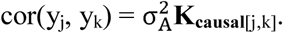

**K**_**causal**_ is an idealized genetic relationship matrix (GRM) that reflects genetic relationships between individuals at an unknown set of causal variants.

Because the set of causal variants is unknown, **K**_**causal**_ has been approximated in the classical literature by the expected relatedness in the pedigree matrix **K**_**PED**_, which equals twice the kinship matrix **Φ**

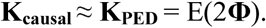

The entries in the kinship matrix **Φ** are known as kinship coefficients. Kinship coefficient φ is the probability that a random allele from subject j is identical by descent (IBD) to an allele at the same locus from subject k. As an example, for a pair of full siblings j and k, this expected probability equals ¼ and therefore **K**_**PED**[j,k]_ = ½. For a design that focuses exclusively on sib pairs and assuming no dominance contribution, an estimate of the additive genetic variance 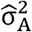 is therefore two times the phenotypic correlation *r*_*P*_ of sib pairs.

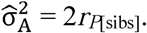

When extended pedigrees are available, the entire **K**_**PED**_ can be leveraged in a linear mixed model (LMM) framework. In this case, the phenotype vector **y** is modelled as

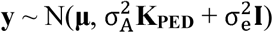

and the genetic variance 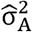 is estimated with restricted maximum likelihood (REML).^6–9^

When genetic data are available, the total IBD fraction of the genome **K**_**IBD**[j,k]_, also referred to as 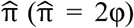, can be estimated for j and k and used in the above LMM framework instead of its expected value from **K**_**PED**[j,k]_. In other words, instead of being approximated with **K**_**PED**_, **K**_**causal**_ is approximated with **K**_**IBD**_.^10,11^

The advent of SNP chips resulted in the use of thousands of markers in the computation of the GRM, thus allowing to estimate heritability in samples without pedigree information.^5,12^ In this case, assuming an N × M genotype matrix 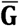 with zero-column mean and unit-column variance, **K**_**causal**_ is approximated with an identity-by-state (IBS) genetic covariance matrix **K**_**IBS**_

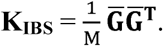

Given that the set of M typed SNPs typically does not include all causal variants and/or it includes tag SNPs that are in imperfect linkage disequilibrium (LD) with said variants, the use of **K**_**IBS**_ can lead to an underestimation of the total heritability. For unrelated individuals, this estimate reflects only the proportion of phenotypic variance captured directly or indirectly by the typed SNPs – i.e. the so-called SNP heritability 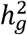 (note that 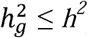). The GCTA software^13^ uses **K**_**IBS**_ on unrelated individuals in order to estimate 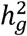. It has been shown that SNP heritability 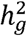 can vary with minor allele frequency (MAF), LD and genotype certainty.^14^ The LDAK software^15^ can be used in order to accommodate these parameters in the computation of **K**_**IBS**_.

More recently, it has been shown that **K**_**IBD**_ can be effectively substituted in the LMM by a truncated IBS genetic covariance matrix **K**_**IBS>t**_, in which all **K**_**IBS**_ entries below a threshold t are set to zero.^16^ Moreover, the same authors introduced a method for the simultaneous estimation of SNP 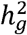 and total heritability *h*^*2*^ by jointly fitting an LMM with **K**_**IBD**_ & **K**_**IBS**_ (or **K**_**IBS>t**_ instead of **K**_**IBD**_). This way, the authors provided heritability estimates with narrower confidence intervals and showed that total heritability estimates under this approach were very similar to those under **K**_**IBD**_ (or **K**_**IBS>t**_) alone.

In order for existing methods to produce meaningful heritability estimates, no population structure should be present in the studied samples.^4^ Population structure can arise when individuals of different ancestry are found in the same sample and/or when individuals are admixed. Individuals from different populations tend to have different minor allele frequencies as well as different environmental exposures.^4^ Because population structure correlates with environmental structure, it can inflate heritability estimates. It has been shown that, for SNP heritability 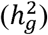 estimates, inclusion of principal components (PCs) as fixed effects cannot fully account for the structure bias.^17^ Moreover, the differences in ancestral allele frequencies affect the computation of **K**_**IBS**_, which ceases to be proportional to **K**_**IBD**_ – even for close relationships. To date, there is not clearly a best strategy for estimating and interpreting heritability in structured/admixed populations.

Nevertheless, an LMM that uses a relationship matrix **K**γ based on local ancestry instead of genotypes has been proposed.^18^ The authors of this method produced accurate estimates of total heritability *h*^*2*^ for several phenotypes in admixed African American samples by fitting an LMM with **K**_γ_ and rescaling its regression coefficient accordingly. However, this method has limitations, as it relies on knowledge about local ancestry and assumes that samples are unrelated.

In light of the above, finding an efficient framework for total narrow-sense heritability estimates in admixed populations with high levels of relatedness remains an open question. Many experimental designs could benefit from a better understanding of how heritability estimates are affected by the simultaneous presence of population and family structure. In this work, we use extensive simulations to evaluate the performance of existing classical and LMM frameworks in estimating SNP 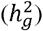 and total (*h*^*2*^) heritability in two population-based cohorts from Greenland. The Greenlanders are an isolate with many unique characteristics, such as small population size, high degree of relatedness and extensive population structure with ancestry from both the Inuit and Europeans.^19,20^ Our goal is to find a way to estimate and understand narrow-sense heritability in such populations, thus gaining valuable insights into the genetic architecture of complex traits in such populations.

## Subjects and Methods

### Samples

#### Greenlanders

The Greenlandic subjects (N = 4,659) came from two general health surveys. The first survey^21^ consisted of Greenlanders living in Denmark (the BHH cohort; N = 546), recruited during 1998-1999, as well as Greenlanders living in Greenland (the B99 cohort; N = 1,328), recruited during 1999-2001 as part of a general population health survey. The second survey consisted of Greenlanders living in Greenland^22^ (the IHIT cohort; N = 2,785), recruited during 2005-2010 as part of a population health survey.

#### Danes

The population-based Danish sample (N = 5,470) was obtained from Inter99,^23^ a randomized intervention study collected at the Research Centre for Prevention and Health. In addition, 513 (N = 1,169) Danish sib pairs were identified across several available Danish cohorts, namely the population-based cohorts (i) Inter99^24^ (N = 294); (ii) Helbred2006^25^ (N = 121); (iii) Helbred2008^26^ (N = 38); and (iv) Helbred2010^27^ (N = 57), all recruited from the Research Centre for Prevention and Health, Glostrup Hospital, Denmark, as well as cohorts collected for the study of type 2 diabetes, namely (v) Vejle Diabetes Biobank^28^ (N = 570), recruited at the Vejle Hospital, Denmark; (v) the ADDITION study^29^ (N = 8), recruited at the Department of General Practice at the University of Aarhus, Denmark; and (vi) SDC^30^ (N = 81) recruited at the outpatient clinic at Steno Diabetes Center, Denmark.

### Ethics statement

The two Greenlandic surveys received ethics approval from the Commission for Scientific Research in Greenland (project 2011-13, ref.no. 2011-056978; project 2013-13, ref.no. 2013-090702), and the Danish studies were approved by the local ethics committees (protocol ref.no. H-3-2012-155). All studies were conducted in compliance with the Helsinki Declaration II and all participants gave their written consent after being informed about the study orally and in writing.

### Genotyping and quality control

Both the Greenlandic and the unrelated Danish samples were typed on Illumina’s Cardio-MetaboChip (Illumina, San Diego, CA, USA). The Cardio-MetaboChip includes 196,725 SNPs selected from genetic studies of cardiovascular, metabolic and anthropometric traits.^31^ Moreover, the unrelated Danish samples and the Danish sib pairs were typed on Illumina’s Infinium OmniExpress chip, which includes ∼710,000 markers. Standard quality control was carried out with PLINK v1.9^32^ and included filtering for *per-individual* (`--mind 0.01`) and *per-marker* (`--geno 0.01`) genotype missingness = 1%. The datasets passing quality control consisted of (i) 4,659 Greenlanders typed on 187,181 Cardio-MetaboChip autosomal SNPs; (ii) 5,470 unrelated Danes typed on 186,639 Cardio-MetaboChip and 618,037 OmniExpress autosomal SNPs; and (iii) 1,169 Danes forming sib pairs typed on 609,605 OmniExpress autosomal SNPs.

### Phenotype simulations

We simulated 1,000 quantitative phenotypes with true total narrow-sense heritability *h*^*2*^ = {0.4, 0.6, 0.8} using real genetic data from 4,659 Greenlanders and 5,470 + 1,169 Danes. We used either GCTA or our own scripts. Simulations were carried out as follows:

First, we defined an N × C causal genotype matrix **G_causal_** by sampling C = 1,500 SNPs from a list of all available SNPs.

Second, we sampled SNP effects, represented by a C × 1 effect vector **b**, from a standard normal distribution N(0,1). In order to model the relationship between the effect size of SNP i and its allele frequency *f*_*i*_, a genotype matrix **G** can be standardized according to the formula

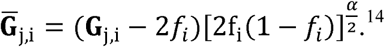

GCTA assumes a specific inverse relationship between SNP effect size and allele frequency by converting the original matrix **G_causal_** into a zero column-mean and unit column-variance standard score matrix 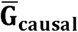. This can be seen as a special case of the above standardizing formula for *α* = - 1.

Third, we computed a vector of polygenic scores **S** by multiplying 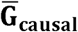 by the corresponding effect vector **b**

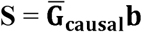

and its additive genetic variance as

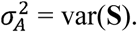

Finally, we computed the phenotype vector **P** by adding an environmental vector ε of i.i.d. error terms to the polygenic score vector **S**

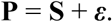

Error terms were sampled from the distribution 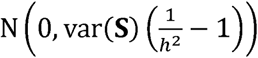.

In some simulations involving the Greenlanders, we also modelled the interaction between environment and ancestry by adding an interaction vector **E×Anc** to the sum

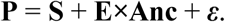

In particular, if 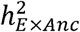 is the proportion of phenotypic variance explained by the interaction and **Q_Inuit_** is the vector of proportion of Inuit ancestry,

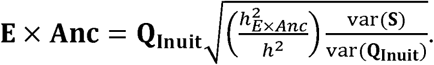

As a consequence, the noise terms in ε are now sampled from 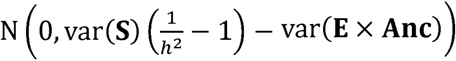.

### Linear mixed model

In an LMM, the phenotype **y** is modelled as a mixture of fixed and random effects (i.e. the effects of the causal variants)

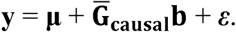

Assuming that 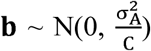 and 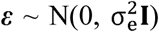 under the GCTA model with *α* = −1, **y** follows a multivariate normal distribution with mean μ and variance

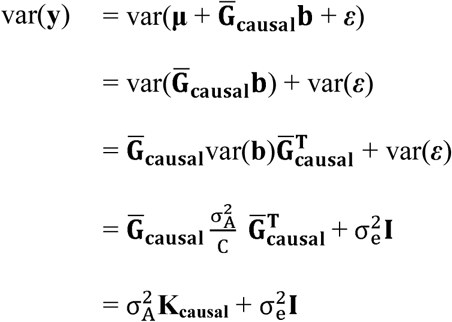

such that

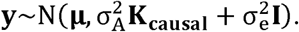

Total narrow-sense heritability is then defined as

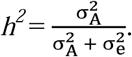

Because **K**_**causal**_ is unknown, we used other GRMs instead, such as **K**_**IBD**_, **K**_**IBS**_, **K**_**IBS>t**_ etc.

### Genetic relationship matrices

We computed the **K**_**IBD**_ matrix for the entire Greenlandic sample from pairwise kinship coefficients 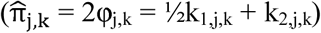 using RelateAdmix^33^ (or alternatively REAP^34^) on a genotype file with MAF cutoff = 0.01. We subsequently identified 1,465 Greenlandic sib pairs by use of empirical thresholding over k_1_ (IBD1; 0.3 < k_1_ ≤ 0.7) and k_2_ (IBD2; 0.1 < k_2_ ≤ 0.5) on the RelateAdmix output. We then re-computed the **K**_**IBD**_ matrix for the identified sib pairs using RelateAdmix. **K**_**IBD**_ for the Danish sib pairs was computed from the PLINK `--genome` output, using MAF cutoff = 0.01. For both the Greenlandic and Danish sib pairs, we also computed a truncated **K**_**IBD>t**_ matrix, in which all entries below a threshold t = 0.05 were set to zero, and a 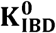 matrix, in which all between-sib pair values were set to zero.

**K**_**IBS**_ and **K**_**IBS>t**_ (t = 0.05) for both Greenlanders and Danes were computed directly with GCTA using a MAF cutoff = 0.01. Causal variants were removed from the computation of **K**_**IBS**_ and **K**_**IBS>t**_. Because causal variants are selected from the entire list of available SNPs, we assume that they have a similar allele frequency distribution as the genotyped SNPs. We can explicitly control for this by adding `--grm-adj 0` to the GCTA command line, however, this setting had no effect on our estimates (data not shown). For some heritability estimations, we computed 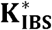 after removing not only the causal variants, but also all variants in LD with the causal ones (i.e. by applying extreme LD pruning in the vicinity of causal variants). Finally, we also estimated heritability by use of a 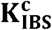 matrix in which all causal variants were included.

### Heritability estimation

Additive genetic variance 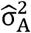 and, subsequently, total narrow-sense heritability *h*^*2*^ were estimated with the GRM-based restricted maximum likelihood (GREML) procedure implemented in GCTA and in LDAK for various GRMs. For the sib pairs in particular, we carried out total narrow-sense heritability estimations using (i) the IBD-based matrices (**K**_**IBD**_, **K**_**IBS>t**_ and 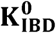) alone; (ii) the IBS-based matrices (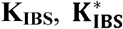 and 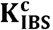) alone; (iii) **K**_**IBS**_ together with **K**_**IBD**_, **K**_**IBS>t**_, 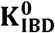 or **K**_**IBS>t**_; and (iv) the classical sib-pair approach. We also estimated 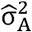 after adjusting for the first 5, 10 or 20 PCs, or proportion of Inuit ancestry (where applicable).

### Analysis settings

We ran phenotype simulations and heritability estimates on three groups of Greenlanders: (i) all samples (N = 4,659); (ii) sib pairs (N = 1,688); and (iii) unrelated individuals (N = 585), as well as two separate groups of Danes: (i) unrelated individuals (N = 5,470); and (ii) sib pairs (N = 1,169). In both populations, the 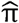 threshold for identifying unrelated individuals was 0.0625. We estimated total heritability on the above groups using different GRMs, without and with covariates (summarized in Table 1). For the sake of simplicity, we do not show results that are nonsensical (e.g. **K**_**IBD**_ for unrelated individuals).

**Table 1:** overview of the different approaches for estimating heritability from simulated data. The linear mixed models (LMMs) were run with and without covariates. Only the meaningful combinations of methods and sample groups were considered

|  |  | Unrelated | Sib pairs | All* |
| --- | --- | --- | --- | --- |
| LMM | $\mathbf{K}_{IBD}$ | N/A | $h^2$ | N/A |
| | $\mathbf{K}_{IBS} / \mathbf{K}_{IBS}^*$ | $h_g^2$ | $h^2$ | N/A |
| | $\mathbf{K}_{IBD} \& \mathbf{K}_{IBS}$ | N/A | $h^2$ and $h_g^2$ | $h^2$ and $h_g^2$ |
| | $\mathbf{K}_{IBS>t} \& \mathbf{K}_{IBS}$ | N/A | $h^2$ and $h_g^2$ | $h^2$ and $h_g^2$ |
| Classical sib-pair analysis | | N/A | $h^2$ | N/A |

\*analysis carried out in Greenlanders only; N/A: not applicable

### Application to real data

We applied the best performing model to real phenotypic data from the two Greenlandic cohorts. All traits considered were quantitative and consisted of basic anthropometric traits (height, weight, body mass index, hip circumference, waist circumference and waist-to-hip ratio), as well as serum lipid levels (total cholesterol, HDL cholesterol, LDL cholesterol and triglycerides). Data were rank-transformed to the quantiles of a standard normal distribution. Age and sex were included as covariates. We also carried out an empirical investigation of the impact of allele frequency and LD weighting – as defined in the LDAK model^14^ – on heritability estimates. In particular, we estimated total narrow-sense heritability for the ten available traits assuming seven different genotype standardizations by setting LDAK’s parameter *α* = {-1.25, −1, −0.75, −0.5, −0.25, 0, 0.25}, and accounting for LD weighting. In this context, the model used by GCTA can be seen as a special case of the LDAK model by setting *α* = −1 and ignoring LD weighting.

## Results

### Admixture and relatedness in the Greenlandic and Danish data

Principal component analysis (PCA) and ancestry component analysis with ADMIXTURE^35^ of 4,659 individuals showed that the general Greenlandic population is the result of admixture between Greenlandic Inuit and European populations and that there is high variance in the admixture profiles of the Greenlanders (Figure 1A).^19,20^ Conversely, a sample of 5,470 Danish individuals appeared largely unstructured (Figure 1B).^36^ We identified a large number of sib pairs in the Greenlandic sample (1,465 pairs; N = 1,688 individuals; Figure S1A). We also confirmed the lack of relatedness in the unrelated Danish samples (Figure S1B) and the presence thereof in the Danish sib pairs (Figure S1C).

**Figure 1:**
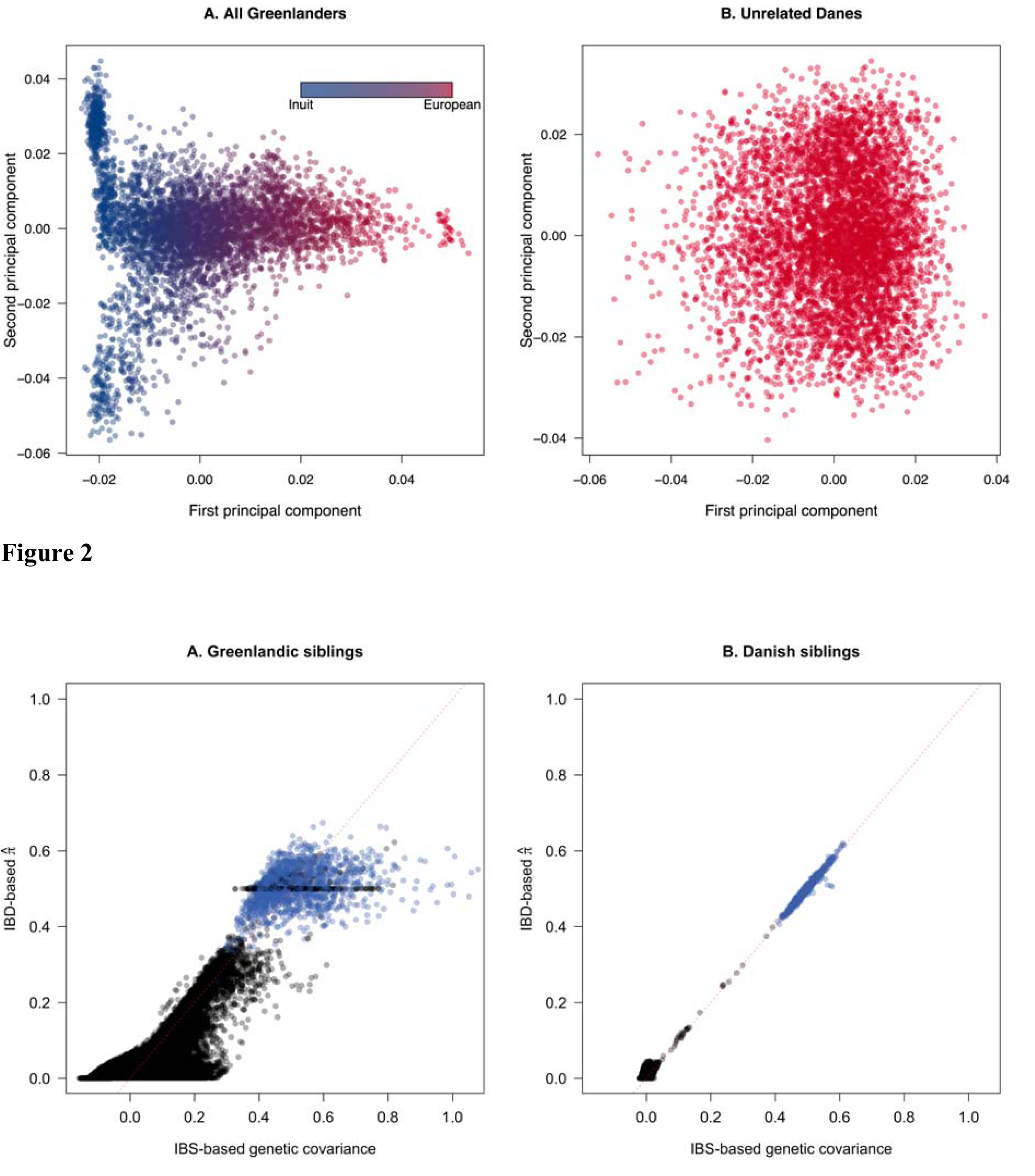
principal component analysis of (A) the entire Greenlandic (N = 4,659) and (B) unrelated Danish (N = 5,470) cohorts. Greenlandic individuals were colored according to their admixture proportions as estimated with ADMIXTURE assuming K = 2 ancestral components. Note that Greenlanders with high proportion of Inuit ancestry (blue points) present further population structure along the second principal component. Due to persisting batch effects, the Danish data were pruned with PLINK (window size = 50; step size = 5; *r*^*2*^ = 0.1) before the analysis.

### Identity by state in the Greenlandic and Danish data

We illustrate the intrinsic differences of IBS between admixed and unadmixed populations by plotting the IBS-based genetic covariance against the IBD-based 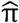 estimates from the Greenlandic and Danish sib pairs, respectively (Figure 2). For a given kinship (e.g. full siblings), the corresponding IBS values were far more dispersed in the Greenlanders (Figure 2A) than in the Danes (Figure 2B). This is due to the heterogeneous admixture profiles in the Greenlanders (Figure 1A). In other words, whereas IBS is proportional to IBD in unadmixed related individuals, this does not hold for the admixed individuals.

**Figure 2:**
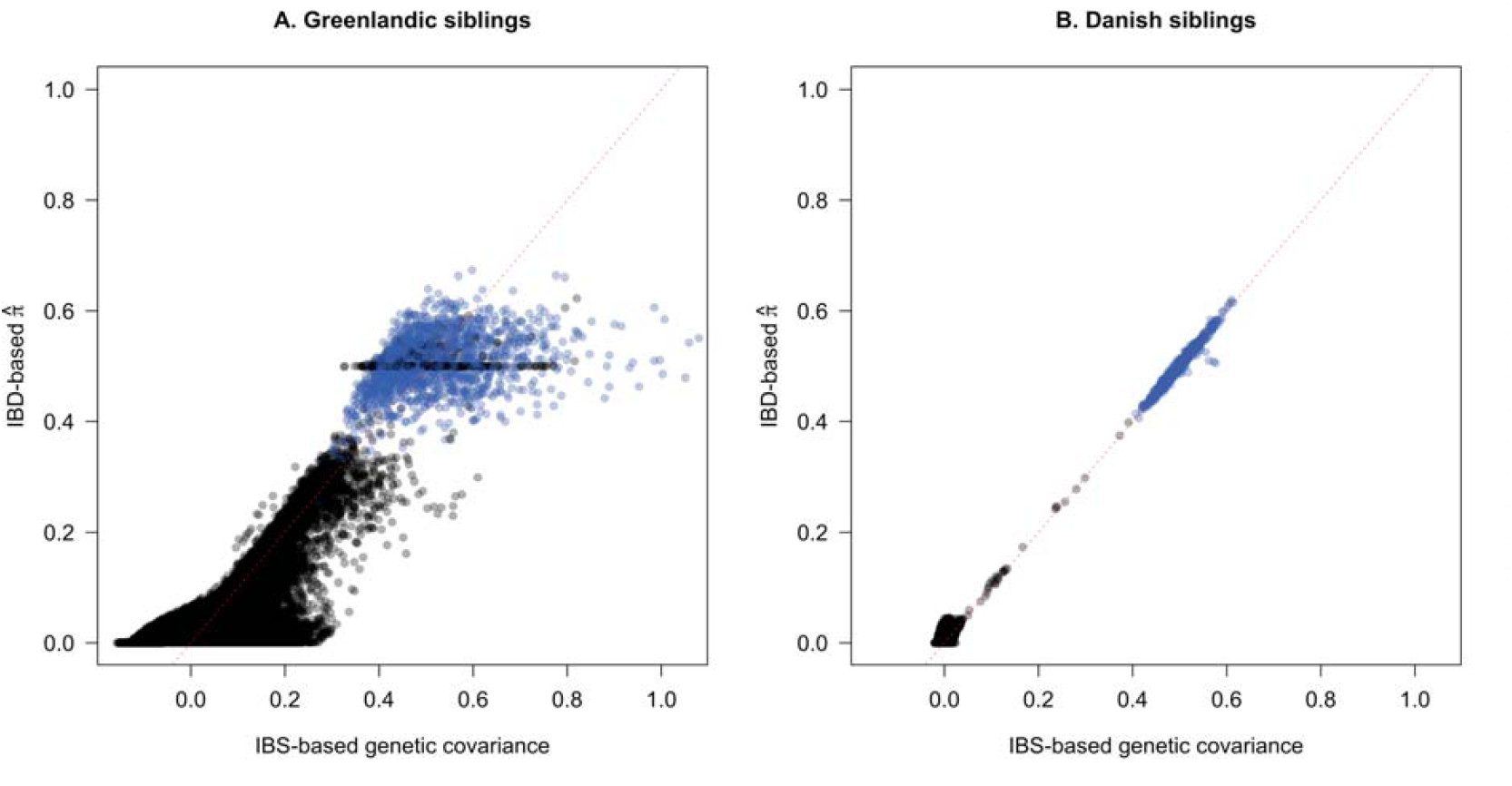
scatterplot of within-pair (blue) and between-pair (black) IBS-based genetic covariance against IBD-based 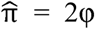 estimates for the (A) Greenlandic (MetaboChip) and (B) Danish (OmniExpress) siblings. IBS values were computed with GCTA, whereas IBD values were computed with RelateAdmix for the Greenlanders and with PLINK for the Danes. A negative IBS value denotes that two individuals are less than average related to each other. An y = x dotted line is shown in red.

### Heritability estimates in phenotypes with no population-specific environmental effects

We explored a number of approaches for estimating narrow-sense heritability *h*^*2*^ in the admixed Greenlandic and the unadmixed Danish population. We simulated quantitative traits by (i) randomly selecting 1,500 causal loci with effect sizes depending on the allele frequency, such that the effect sizes of the standardized genotypes are normally distributed as assumed in the GCTA software, and (ii) adding environmental noise so that the true simulated *h*^*2*^ was 0.4, 0.6 or 0.8. In these simulations, all individuals were set to having the same environmental variance regardless of their ancestry. We then estimated *h*^*2*^ in an LMM framework for different GRMs – namely **K**_**IBD**_, **K**_**IBS**_, **K**_**IBS>t**_ and combinations thereof (Figure 3; Figure S2; Figure S3).

**Figure 3:**
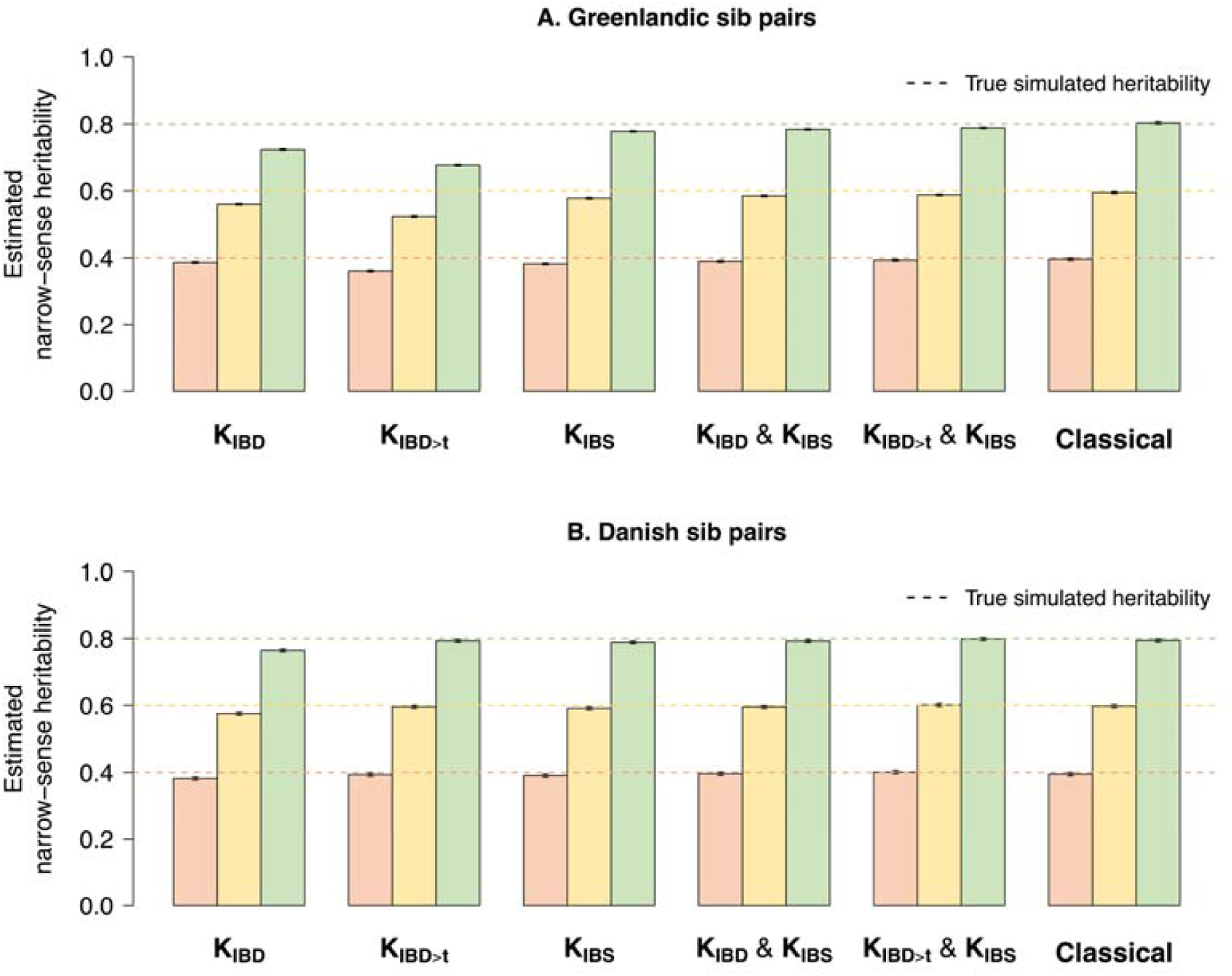
mean total heritability estimates from 1,000 simulated phenotypes and 95% confidence intervals of the sampling distribution in (A) Greenlandic and (B) Danish sib pairs under various LMMs and the classical approach with no ancestry-specific environmental interactions added to the phenotypes. Genotype scaling parameter *α* was set to −1 (GCTA’s standard) for both phenotype simulation and heritability estimation. **K**_**IBD**_: IBD-based GRM; **K**_**IBD>t**_: IBD-based GRM in which all entries below t = 0.05 were set to zero; **K**_**IBS**_: IBS-based GRM.

#### Total heritability estimates in sib pairs using one GRM

The use of **K**_**IBD**_, which captures the fraction of the genome shared IBD 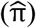, resulted in underestimates of total heritability both in the Greenlandic and the Danish sib pairs (Figure 3). Total heritability in the Greenlandic sib pairs was also underestimated when we used **K**_**IBD>t**_ (Figure 3A) and 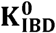 (Figure S2A). Conversely, using **K**_**IBD>t**_ or 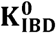 in the Danish sib pairs did not result in any significant downward biases (Figure 3B; Figure S2B). These results were insensitive to the method of IBD inference: both our method of choice (RelateAdmix^33^) and an alternative (REAP^34^) returned similar results (data not shown).

For closely related unadmixed individuals, IBS is proportional to IBD and therefore estimates based on **K**_**IBS**_ will correspond to the total narrow-sense heritability as well.^37^ Indeed, simulated *h*^*2*^ was fully recovered in the Danish sib pairs when **K**_**IBS**_ was used (Figure 3B), but showed consistent downward biases across all simulated *h*^*2*^ values in the Greenlandic sib pairs (Figure 3A). Removing all SNPs with any LD with the causal variants (*r*^*2*^ = 0) from the GRM (i.e. using the 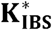 matrix) returned lower yet still comparable *h*^*2*^ estimates in both the Greenlandic (Figure S2A) and Danish sib pairs (Figure S2B). Conversely, when causal variants were included in the computation of the GRM (i.e. using the 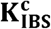 **matrix**), then 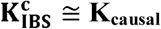 and, consequently, *h*^*2*^ was recovered (Figure S4).

#### Total heritability estimates in sib pairs using two GRMs

The use of two GRMs (e.g. **K**_**IBD**_ & **K**_**IBS**_) for heritability estimates is meant to leverage datasets in which both closely and more distantly related individuals are present.^16^ However, when we applied this approach to the entire Greenlandic dataset (N = 4,659), we observed a downward bias in total heritability estimates for the **K**_**IBD**_ & **K**_**IBS**_ model, while using **K**_**IBS>t**_ & **K**_**IBS**_ erroneously returned heritability estimates near 1.00 regardless of the true simulated value (Figure S5). Nevertheless, when we performed the two-GRM analysis on the 1,465 Greenlandic sib pairs alone, the true simulated *h*^*2*^ was almost perfectly recovered for all GRM combinations, with estimates showing only a minor downward bias (Figure 3A; Figure S2A). Interestingly, these models outperform the classical sib-pair analysis as evidenced by their lower root-mean-square deviation (Table S1; Figure S3). Extending the sample to include more distant relatives 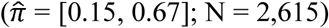 resulted in underestimates of the total heritability (Figure S6), implying that the **K**_**IBD**_ & **K**_**IBS**_ model performs efficiently only on first-degree relatives when admixture is present. We therefore further examined the **K**_**IBD**_ & **K**_**IBS**_ model in the following section, focusing our attention on the sib pairs.

#### SNP heritability estimates in unrelated individuals

The use of **K**_**IBS**_ on unrelated individuals yields SNP 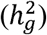 rather than total heritability (*h*^*2*^) estimates.^5^ Bearing this in mind, we also estimated 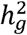 in both unrelated Greenlanders and unrelated Danes (Figure S7; Table S2). In all cases, we found that 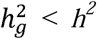. For the Danish samples in particular, the MetaboChip 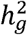 was smaller than the OmniExpress 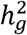.

### Heritability estimates in phenotypes with population-specific environmental effects

In all of the above simulations, we assumed that the environmental component was independent of ancestry. However, when we added an environmental component correlating with ancestry to the simulated phenotypes, the use of **K**_**IBS**_ in the unrelated Greenlanders led to overestimates of SNP heritability 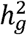, despite adjusting for population structure (Table S2).

Adjusting the **K**_**IBD**_ model for either the first 10 PCs or proportion of Inuit ancestry produced a consistent yet uninterpretable pattern along the 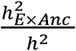 ratio (Figure S8). On the contrary, adjusting the **K**_**IBD**_ & **K**_**IBS**_ model for the same covariates produced a predictable as well as interpretable pattern across all choices for the 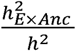 ratio (Figure 4): even though the true simulated heritability 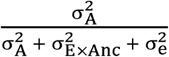 was not recovered, the resulting estimates corresponded to removing the environmental interaction component 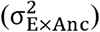 from the denominator of the formula (i.e.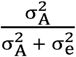) through adjustment for ancestry. We note that adjustment for 10 PCs was equivalent to adjustment for proportion of Inuit ancestry (Figure 4).

**Figure 4:**
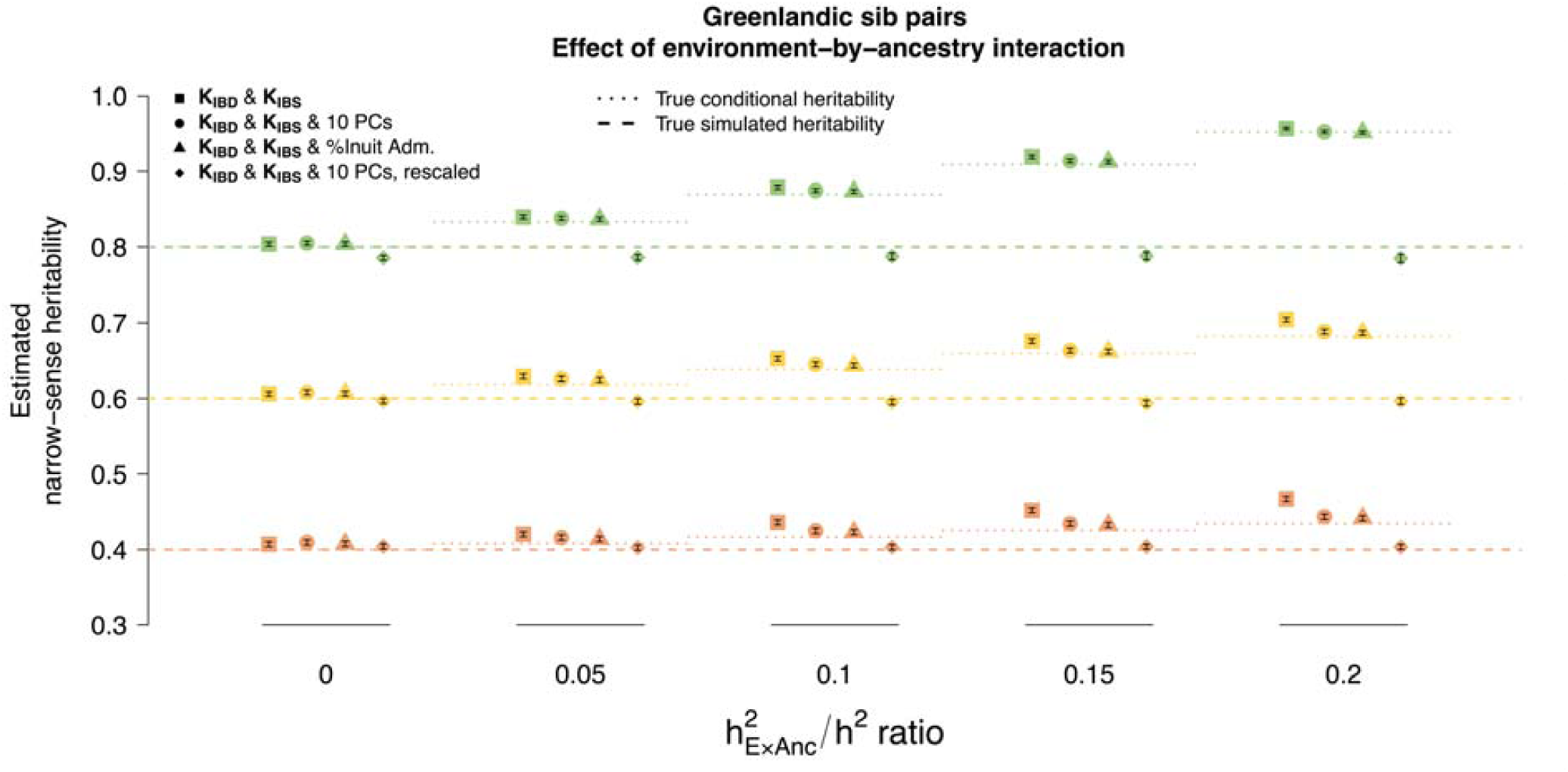
mean total heritability estimates from 1,000 simulated phenotypes and 95% confidence intervals of the sampling distribution in Greenlandic sib pairs under the **K**_**IBD**_ & **K**_**IBS**_ model, without (squares) and with (circles, triangles) population structure covariates, as well as after the additional PCA-based adjustment (diamonds). The relationship between the environment-by-ancestry (ExAnc) interaction and the true simulated heritability is quantified by the 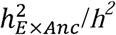 ratio on the x-axis. The models with the covariates (circles and triangles) correspond to estimates of conditional total heritability after adjusting for the ExAnc effect (dotted line). Further rescaling of the conditional heritability estimates by 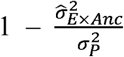 returns the marginal simulated heritability (dashed line).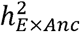: proportion of variance captured by the ExAnc interaction; Inuit adm.: proportion of Inuit admixture. Adjustment for 5 or 20 first PCs returned virtually identical results (data not shown).

Thanks to the interpretability of the resulting “conditional” estimates, we were able to recover the true simulated – the “marginal” – heritability^38^. We achieved this by rescaling the resulting total heritability estimates by a factor of 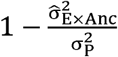, where 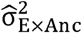 is an estimate of the environmental interaction variance computed as 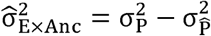. 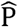 is the phenotype residuals after regressing out the effect of structure captured either by admixture proportions or the first two principal components.

### Application to real phenotypes

We applied the best model (i.e. **K**_**IBD**_ & **K**_**IBS**_ & 10 PCs) and the follow-up PCA-based adjustment to ten quantitative traits in the 1,465 Greenlandic sib pairs (Table 2). Not all phenotypes were equally sensitive to the PCA-based adjustment of their estimated conditional heritability, implying trait-specific environment-by-ancestry interactions. The GCTA model accepts only one type of genotype standardization (*α* = −1), resulting in strong assumptions about the distribution of effect sizes. We therefore also used the LDAK model^14^ and found that the optimal *α* value for genotype standardization varied across traits with most phenotypes supporting *α* ≥ −0.5 (Figure S9; Table S3). Total heritability estimates under the LDAK model^14^ were generally higher than under GCTA (Table 2; Table S3), with the greatest difference observed for height (0.657±0.042 for GCTA against 0.786±0.041 for LDAK). In addition, heritability estimates in eight out of ten real phenotypes were smaller in the Greenlanders than in their European or Mexican counterparts estimated with similar models.

**Table 2:**
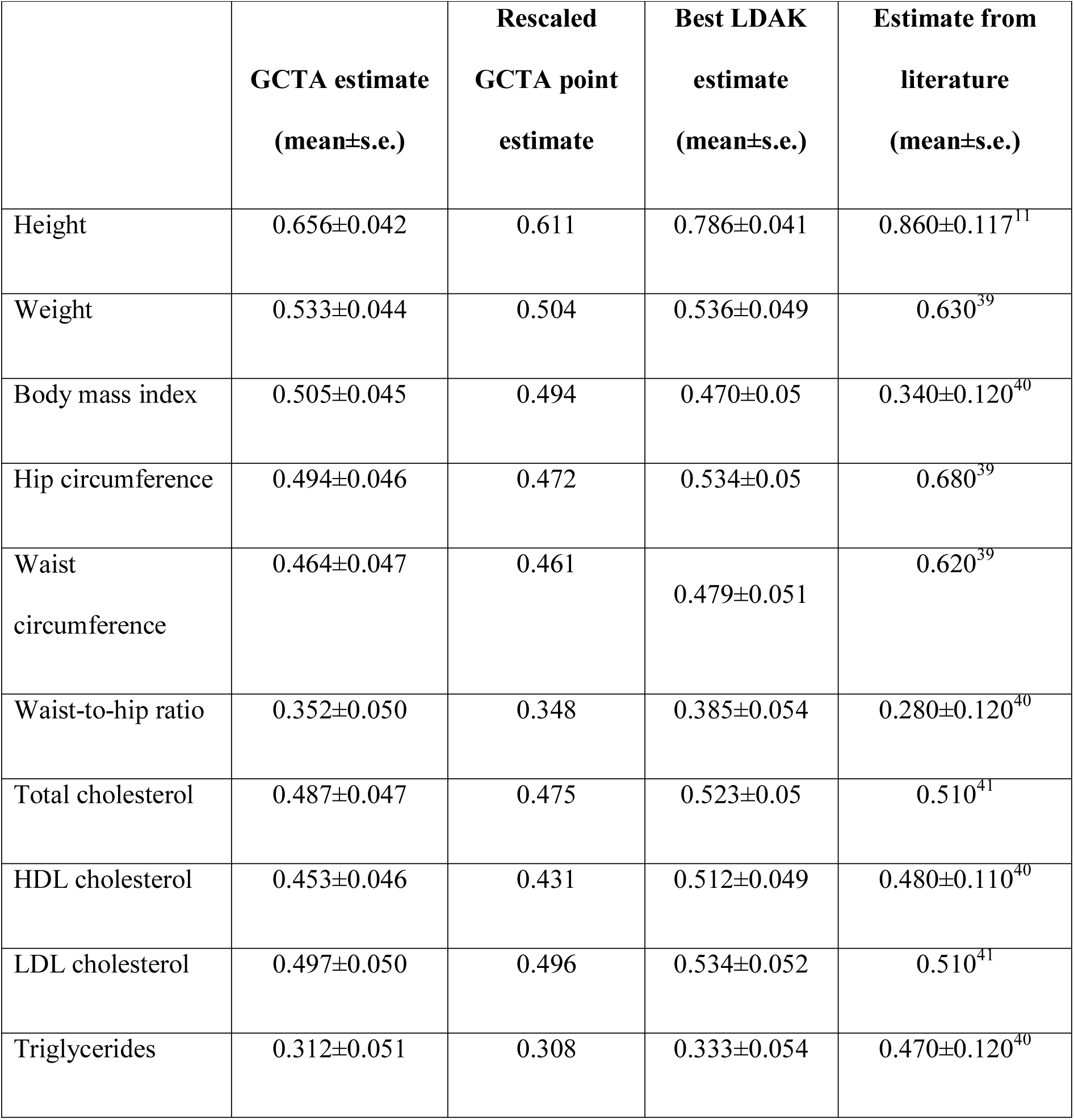
total narrow-sense heritability estimates for real phenotypes under the **K**_**IBD**_ & **K**_**IBS**_ & age & sex & 10 PCs model using (i) the GCTA model; (ii) GCTA followed by PCA-based rescaling; and (iii) the LDAK model with an optimal trait-specific parameter *α* and LD weighting, as well as comparison with (iv) published total heritability estimates in Europeans and Mexicans

|  | **GCTA estimate**<br>**(mean±s.e.)** | **Rescaled**<br>**GCTA point**<br>**estimate** | **Best LDAK**<br>**estimate**<br>**(mean±s.e.)** | **Estimate from**<br>**literature**<br>**(mean±s.e.)** |
| --- | --- | --- | --- | --- |
| Height | 0.656±0.042 | 0.611 | 0.786±0.041 | 0.860±0.117<sup>11</sup> |
| Weight | 0.533±0.044 | 0.504 | 0.536±0.049 | 0.630<sup>39</sup> |
| Body mass index | 0.505±0.045 | 0.494 | 0.470±0.05 | 0.340±0.120<sup>40</sup> |
| Hip circumference | 0.494±0.046 | 0.472 | 0.534±0.05 | 0.680<sup>39</sup> |
| Waist<br>circumference | 0.464±0.047 | 0.461 | 0.479±0.051 | 0.620<sup>39</sup> |
| Waist-to-hip ratio | 0.352±0.050 | 0.348 | 0.385±0.054 | 0.280±0.120<sup>40</sup> |
| Total cholesterol | 0.487±0.047 | 0.475 | 0.523±0.05 | 0.510<sup>41</sup> |
| HDL cholesterol | 0.453±0.046 | 0.431 | 0.512±0.049 | 0.480±0.110<sup>40</sup> |
| LDL cholesterol | 0.497±0.050 | 0.496 | 0.534±0.052 | 0.510<sup>41</sup> |
| Triglycerides | 0.312±0.051 | 0.308 | 0.333±0.054 | 0.470±0.120<sup>40</sup> |

## Discussion

In this work, we explored the performance of existing methods for heritability estimates in the admixed Greenlandic population. Our goal was to propose a framework for unbiased heritability estimates in datasets where both population and family structure are notably present, as well as a way to interpret the resulting estimates. Even though the main focus is on total narrow-sense heritability (*h*^*2*^), we also report results for SNP heritability 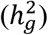, a quantity that has gained a lot of attention in the past decade due to the availability of GWAS data.^5,12,17^

Through extensive simulations, we observed that all LMMs using one GRM led to downward biases in total heritability estimates when applied to family data from Greenland. Common choices of GRM, such as **K**_**IBD**_ and **K**_**IBS**_, led to underestimates of total heritability in Greenlandic sib pairs, whereas no such biases were generally observed for the Danish sib pairs, indicating that admixture exerts a biasing effect on both IBD- and IBS-based estimates. Even though this is not surprising for the IBS-based estimates, as the IBS ∼ IBD assumption does not hold for the Greenlanders, it is not very clear why IBD-based heritability estimates are also affected by admixture.

We also observed that an LMM with two GRMs (**K**_**IBD**_ & **K**_**IBS**_), a method designed to work on data with notable presence of family structure,^16^ also led to downward biases in total heritability estimates when applied to the entire dataset from Greenland. However, when the same analysis focused on the Greenlandic sib pairs, it returned nearly unbiased heritability estimates. This could be due to the fact that, by restricting the analysis to the sib pairs, we controlled more efficiently for the noise that comes from unmatched between-pair admixture proportions. We note that the **K**_**IBD**_ & **K**_**IBS**_ model outperformed the **K**_**IBD**_ model in the Danes, rendering more advisable the use of two GRMs in total narrow sense heritability estimates in unadmixed populations too.

When there is no environmental correlation with ancestry, the **K**_**IBD**_ & **K**_**IBS**_ (or any other combination of one IBD- and one IBS-based GRM) model provides a nearly accurate estimate of the true heritability matched only by the classical sib pair analysis. However, in reality we expect environmental structure, which correlates with genetic structure, to exert an inflating effect on true heritability estimates but found that adjusting for structure did not remove the inflation. Nevertheless, we provide a way to interpret the resulting total heritability estimates from the **K**_**IBD**_ & **K**_**IBS**_ & 10 PCs (or **K**_**IBS>t**_ & **K**_**IBS**_ & 10 PCs) model, as well as a way to adjust for the inflation. In particular, this inflated quantity is referred to as “conditional heritability” in a recent paper,^38^ resulting after adjustment for model covariates like in our case. We observed that, under the **K**_**IBD**_ & **K**_**IBS**_ & 10 PCs (or **K**_**IBS>t**_ & **K**_**IBS**_ & 10 PCs) models, the resulting conditional heritability^38^ estimate will be inflated by a factor of 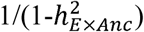 and we propose an adjustment that accounts efficiently for this inflation in order to retrieve the “marginal heritability”.^38^

As for the total narrow-sense heritability estimates of the real phenotypes obtained with the best model (**K**_**IBD**_ & **K**_**IBS**_ & 10 PCs), we observe that in some occasions these are lower for the Greenlandic population than for European populations. A notable example is height, for which total heritability in the Greenlanders was estimated to be 0.656±0.042 (0.611 after the PCA-based adjustment), whereas in unadmixed Europeans it was estimated at 0.860.^11^ We believe that this could be due to the reduced genetic diversity observed in the Greenlanders as a consequence of their particular population history, which included an extreme and prolonged bottleneck in recent times,^19,20^ even though we did observe a notable increase when LD weighting was included in the estimation model according to LDAK.^14^

Finally, our results from SNP heritability 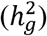 estimates in unrelated Greenlanders were consistent with previous findings, whereby genetic structure was reported to inflate heritability estimates when the environment was not constant;^17^ we found that accounting for structure would not always suffice. In any case, SNP heritability estimates in the Greenlanders should be interpreted with caution because, as we saw, IBS is affected by admixture that can lead to artificially increased levels of LD between causal and typed markers.

In summary, we advise against the use of **K**_**IBD**_ or **K**_**IBS**_ alone for total narrow-sense heritability estimates in populations with substantial levels of population and family structure. Instead, **K**_**IBD**_ & **K**_**IBS**_ & 10 PCs on a subset with high relatedness (preferably sib pairs) is advisable given that **K**_**IBD**_ can now be efficiently computed for admixed populations.^33,34^ As an alternative to **K**_**IBD**_, **K**_**IBS>t**_ can be used together with **K**_**IBS**_.^16^ In any case, the resulting conditional *h*^*2*^ estimates should be viewed as potentially inflated by a factor that we estimated at 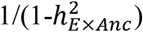 and an additional PCA-based adjustment should be carried out in order to recover the marginal total *h*^*2*^ estimate.

## Supplemental Data

Supplemental Data include 9 figures and 3 tables.

## Acknowledgements

We would like to thank the staff and participants of in the IHIT, B99, and BBH cohorts facilitating this study. The staff and steering committees from Research Centre for Prevention and Health, Glostrup, Denmark, from the ADDITION-DK study, University of Aarhus, Denmark, from Vejle Diabetes Biobank, Vejle Hospital, Denmark, and from Steno Diabetes Center, Gentofte, Denmark are acknowledged for their contribution to collecting and characterizing the Danish cohorts.

---

**Figure S1:**
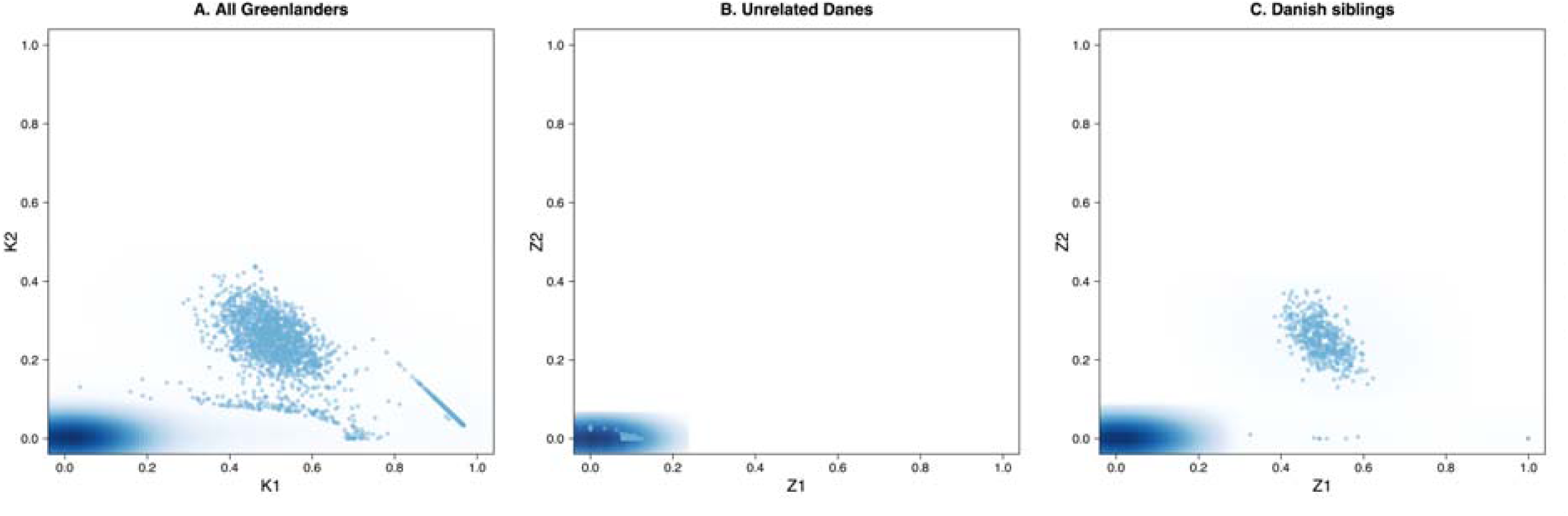
plots of the estimated fraction of the genome where one allele is shared IBD (K1 or Z1) against the estimated fraction where two alleles are shared IBD (K2 or Z2) for (A) all pairs of Greenlandic individuals (Metabochip); (B) unrelated Danish individuals (OmniExpress); (C) and Danish sib pairs (Omniexpress).

**Figure S2:**
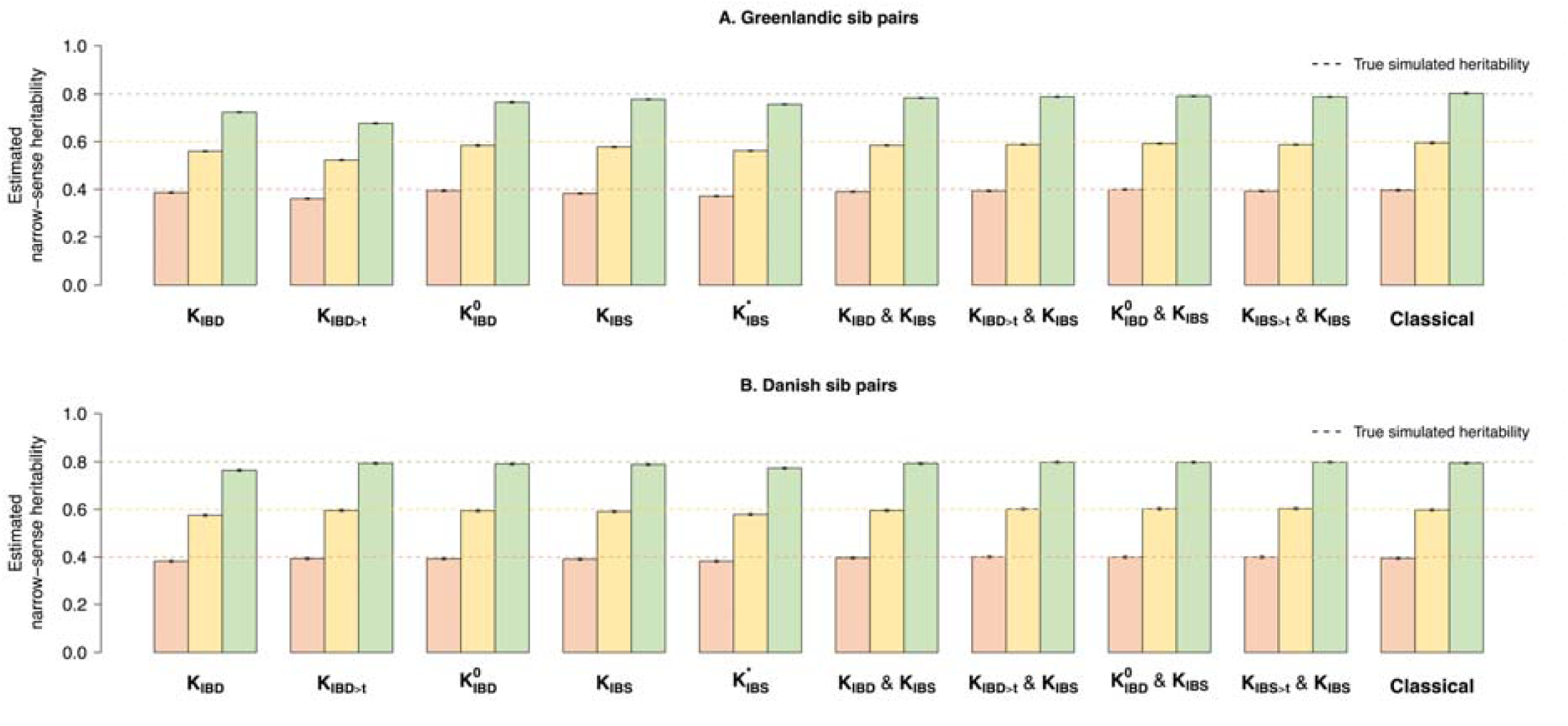
mean total heritability estimates from 1,000 simulated phenotypes and 95% confidence intervals of the sampling distribution in (A) Greenlandic and (B) Danish sib pairs under various LMMs and the classical approach with no ancestry-specific environmental interactions added to the phenotypes. Genotype scaling parameter *α* was set to −1 (GCTA’s standard) for both phenotype simulation and heritability estimation. **K**_**IBD**_: IBD-based GRM; **K**_**IBD>t**_: IBD-based GRM in which all entries below t = 0.05 were set to zero; 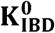: IBD-based GRM in which all between-sib pair entries were set to zero; **K**_**IBS**_: IBS-based GRM; 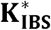: IBS-based GRM in which both causal SNPs and any SNPs in LD with them were omitted; **K**_**IBS>t**_: IBS-based GRM in which all entries below t = 0.05 were set to zero.

**Figure S3:**
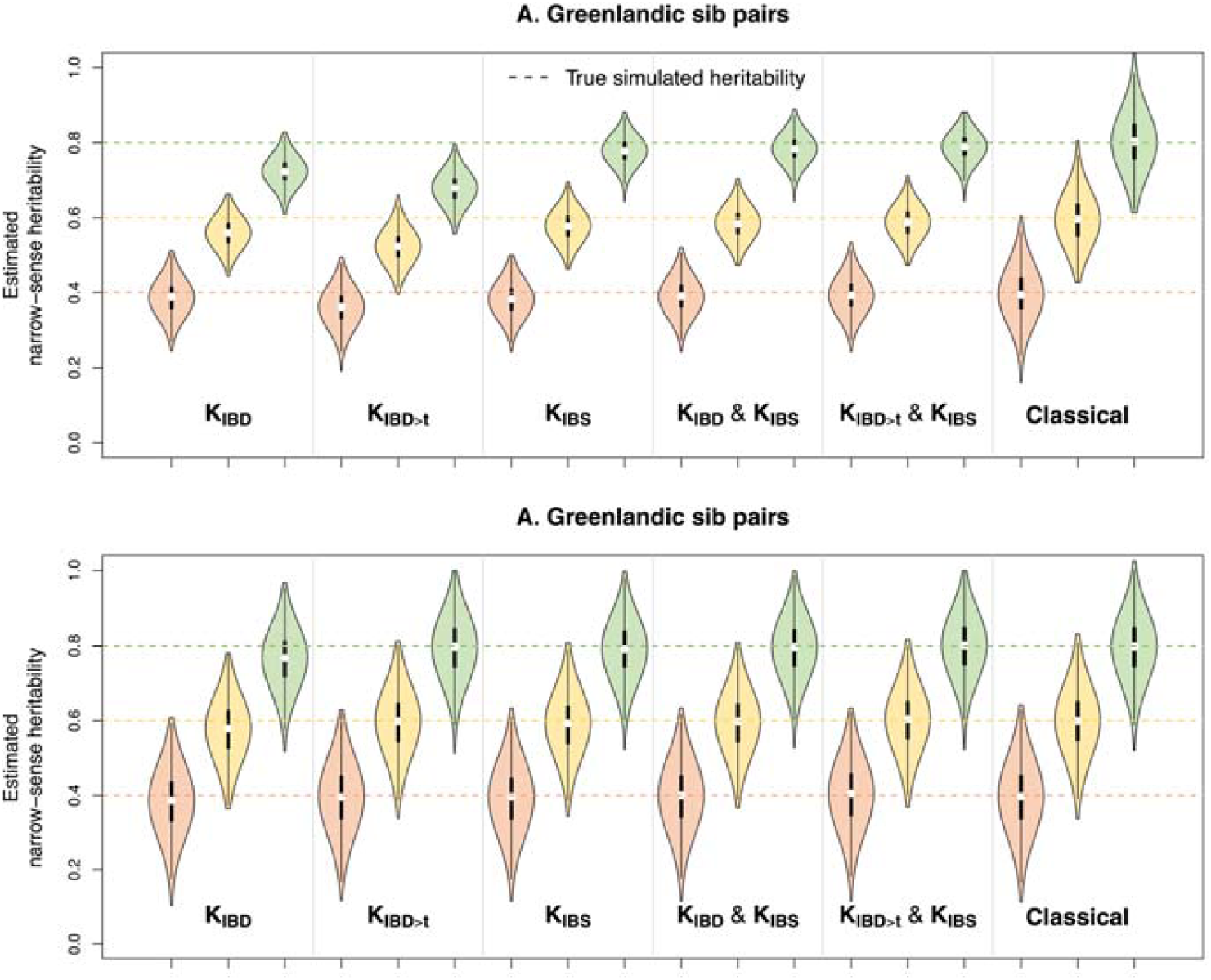
violin plot showing total heritability estimates from 1,000 simulated phenotypes in (A) Greenlandic and (B) Danish sib pairs under various LMMs and the classical approach with no ancestry-specific environmental interactions added to the phenotypes. The median values are shown as white circles, the interquartile ranges as black bars, and the 5^th^ and 95^th^ percentiles as whiskers. Genotype scaling parameter *α* was set to −1 (GCTA’s standard) for both phenotype simulation and heritability estimation.

**Figure S4:**
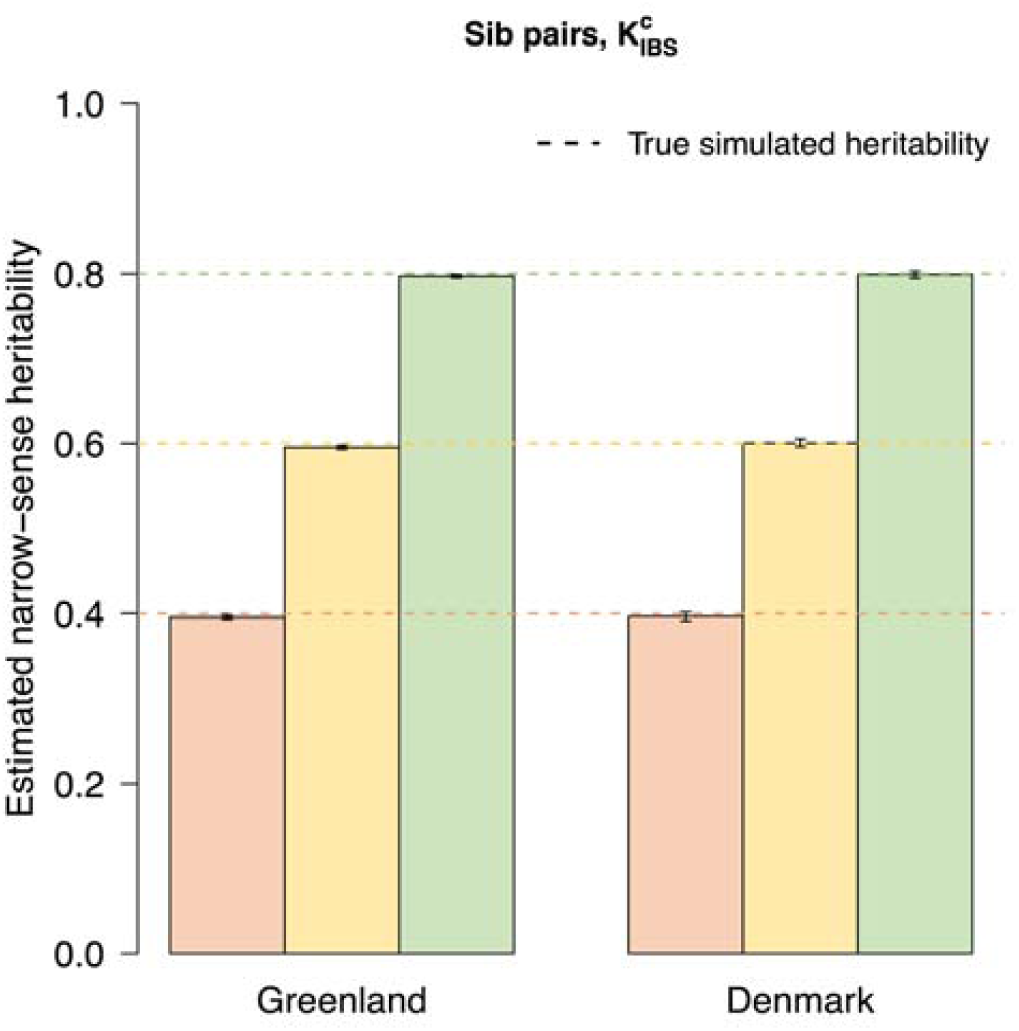
mean total heritability estimates from 1,000 simulated phenotypes and 95% confidence intervals of the sampling distribution in Greenlandic and Danish sib pairs when the IBS-based GRM includes the causal variants 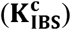.

**Figure S5:**
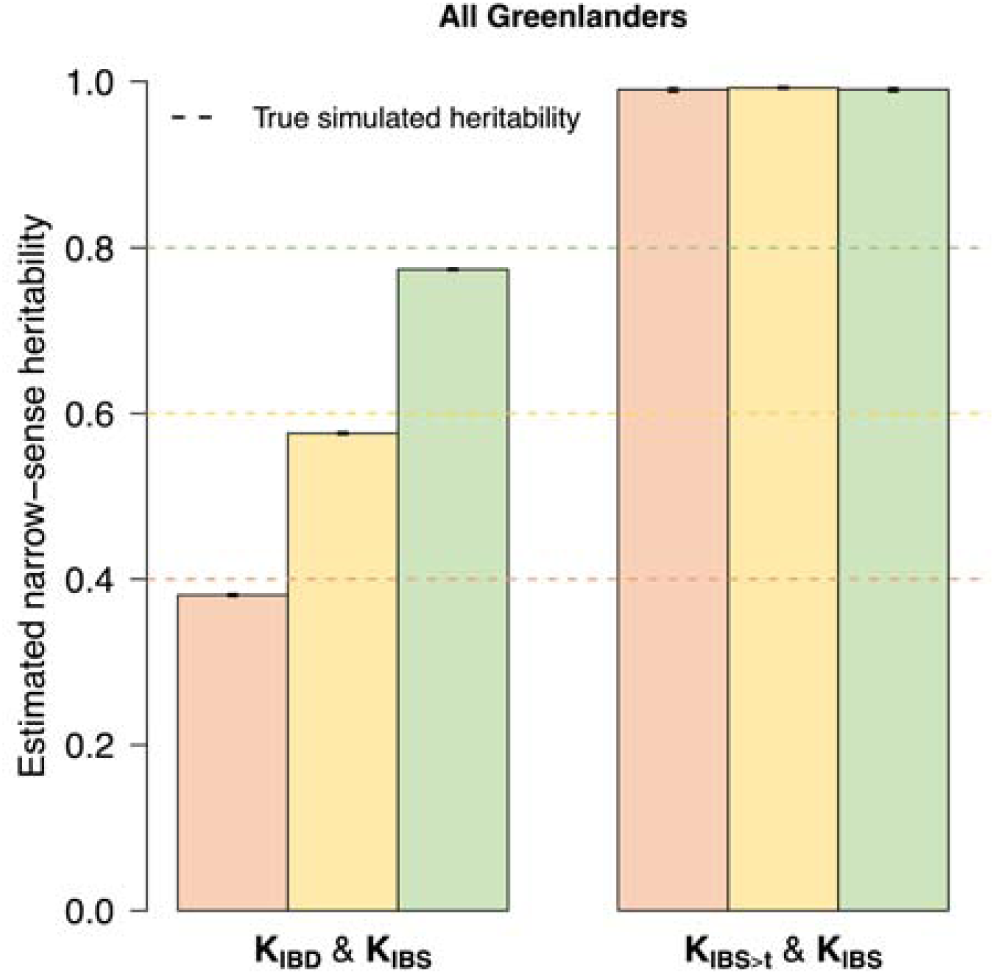
mean total heritability estimates from 1,000 simulated phenotypes and 95% confidence intervals of the sampling distribution in the entire Greenlandic sample under the (A) **K**_**IBD**_ & **K**_**IBS**_ and (B) **K**_**IBS>t**_ & **K**_**IBS**_ models.

**Figure S6:**
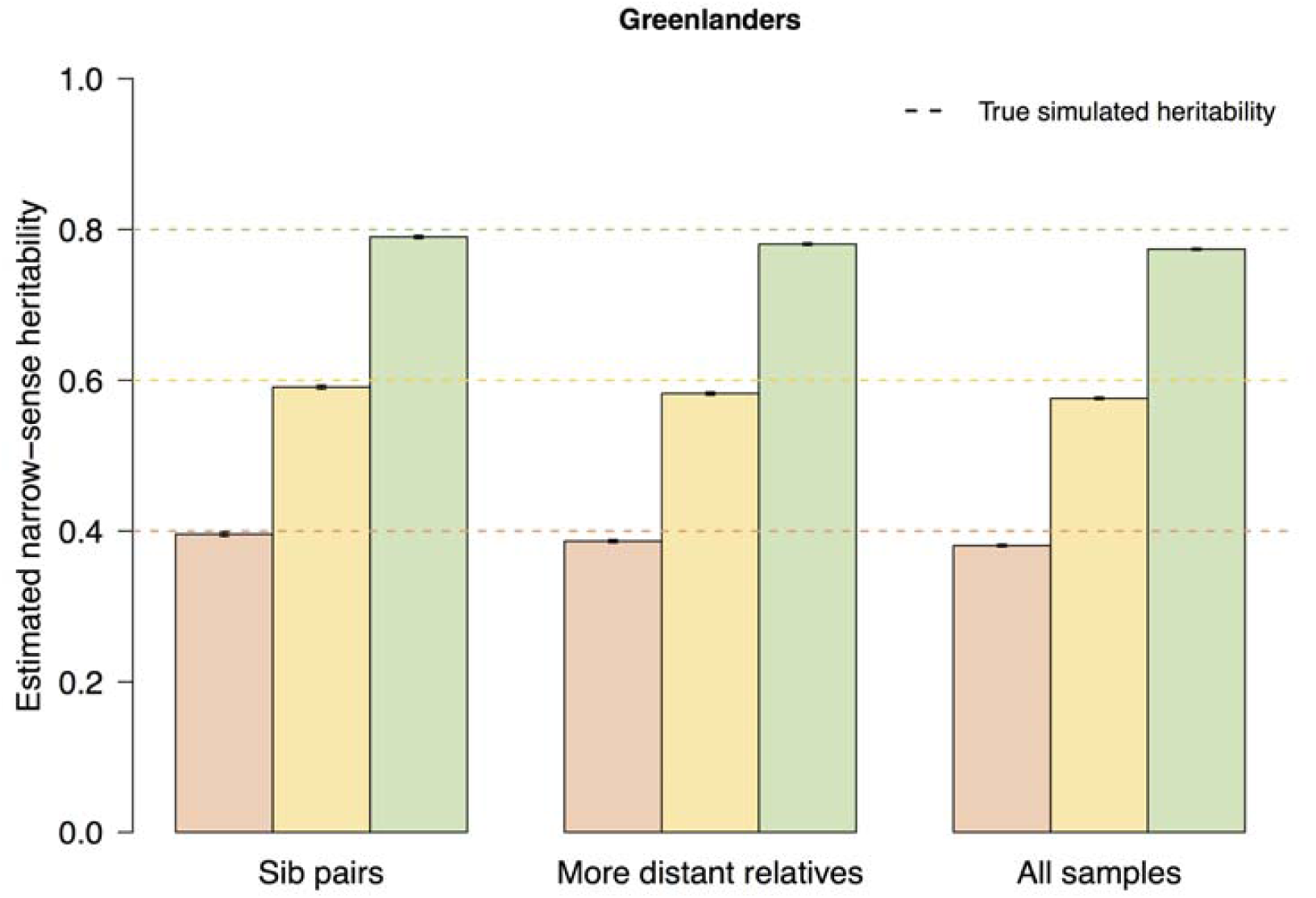
mean total heritability estimates from 1,000 simulated phenotypes and 95% confidence intervals of the sampling distribution under the **K**_**IBD**_ & **K**_**IBS**_ model in Greenlandic sib pairs (left); Greenlandic sib pairs including more distant relatives (middle); and the entire Greenlandic sample (right).

**Figure S7:**
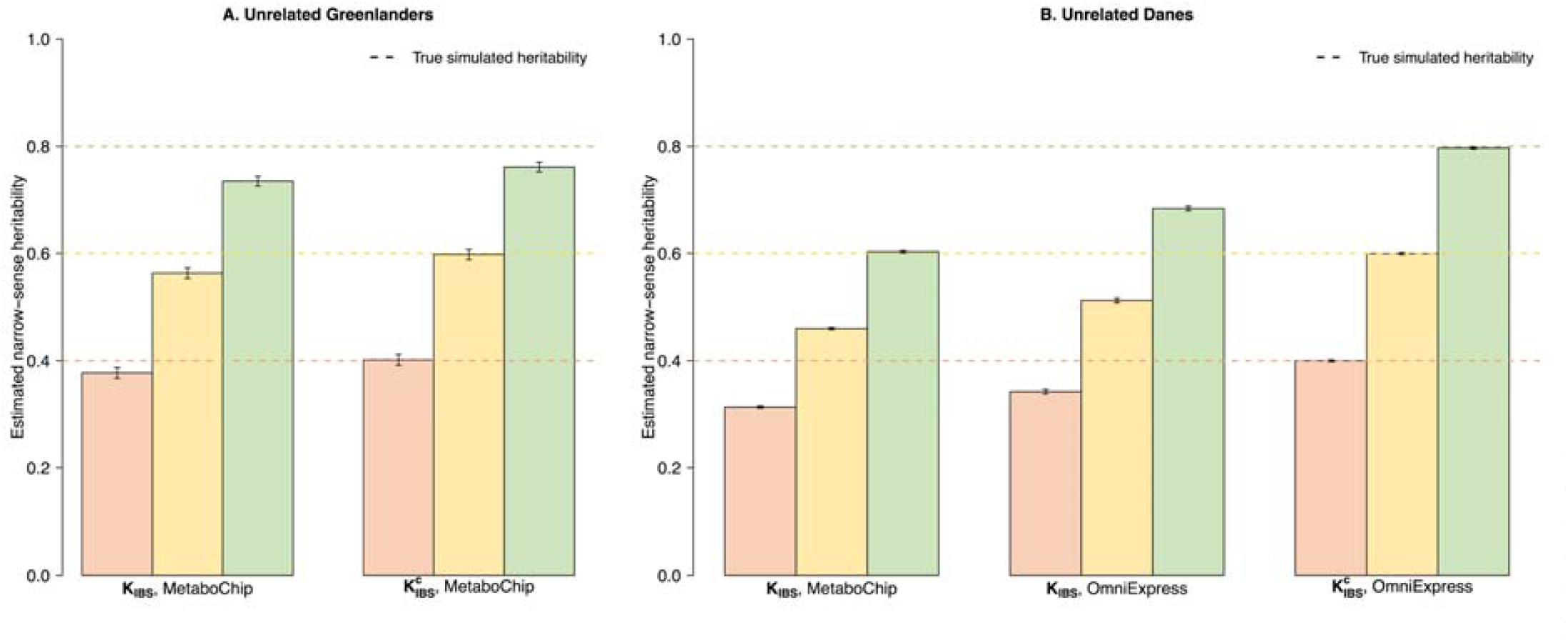
mean IBS-based heritability estimates from 1,000 simulated phenotypes and 95% confidence intervals of the sampling distribution in (A) unrelated Greenlandic and (B) unrelated Danish individuals. Estimations were carried out using both **K**_**IBS**_, returning the SNP heritability, and 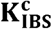, returning the total heritability. SNP heritability estimation in Danes was carried using both the MetaboChip and the OmniExpress data. SNP heritability was lower for the MetaboChip than for the OmniExpress array data, because the MetaboChip SNPs capture a smaller fraction of the genome through LD compared to the denser OmniExpress array. When the causal variants were included in the GRM 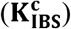, total narrow-sense heritability was almost perfectly recovered.

**Figure S8:**
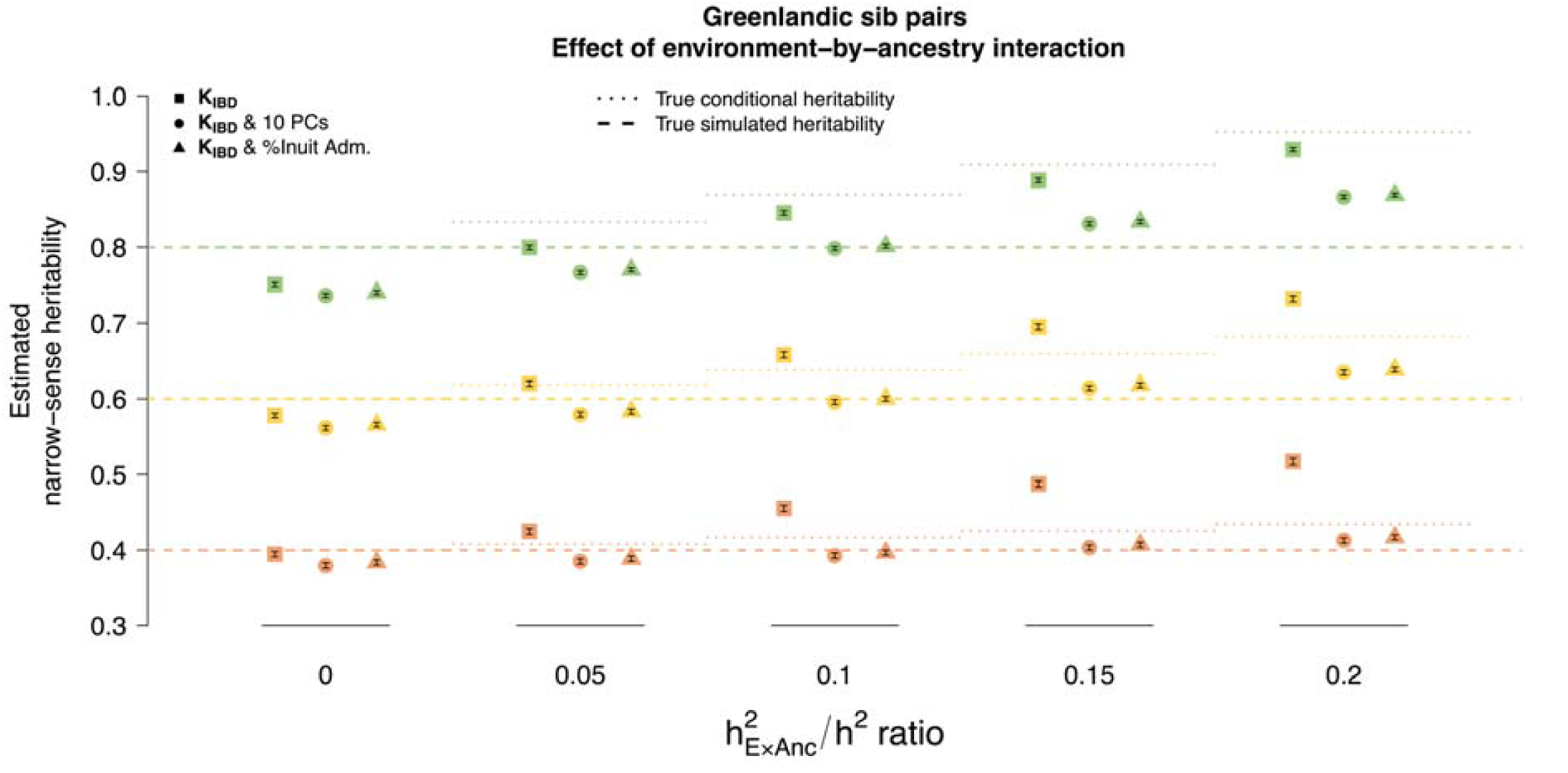
mean total heritability estimates from 1,000 simulated phenotypes and 95% confidence intervals of the sampling distribution in Greenlandic sib pairs under the **K**_**IBD**_ model, without (squares) and with (circles, triangles) population structure covariates. The relationship between the ExAnc interaction and the simulated total heritability is quantified by the 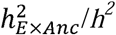 ratio on the x-axis. None of the models recover the true estimated heritability in a systematic and interpretable manner along the 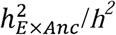 ratio.

**Figure S9:**
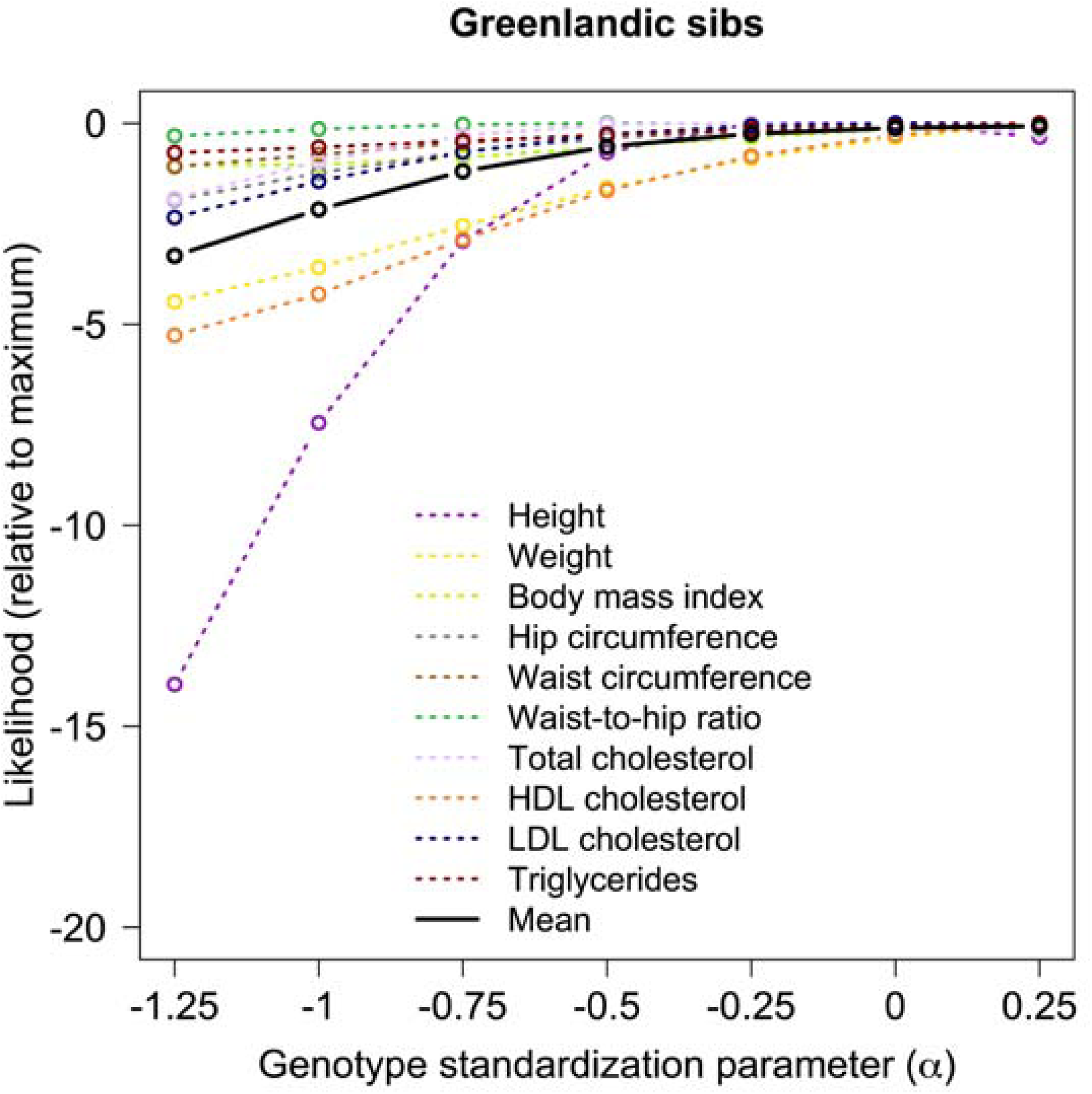
fitness comparisons of the genotype scaling parameter *α* under the LDAK model on the basis of likelihood, whereby higher likelihood indicates better-fitting *α*. Lines report log likelihoods from LDAK for seven values of *α* relative to the highest observed likelihood. Colored lines indicate the ten quantitative traits and the black line reports averages across all traits.

**Table S1:**
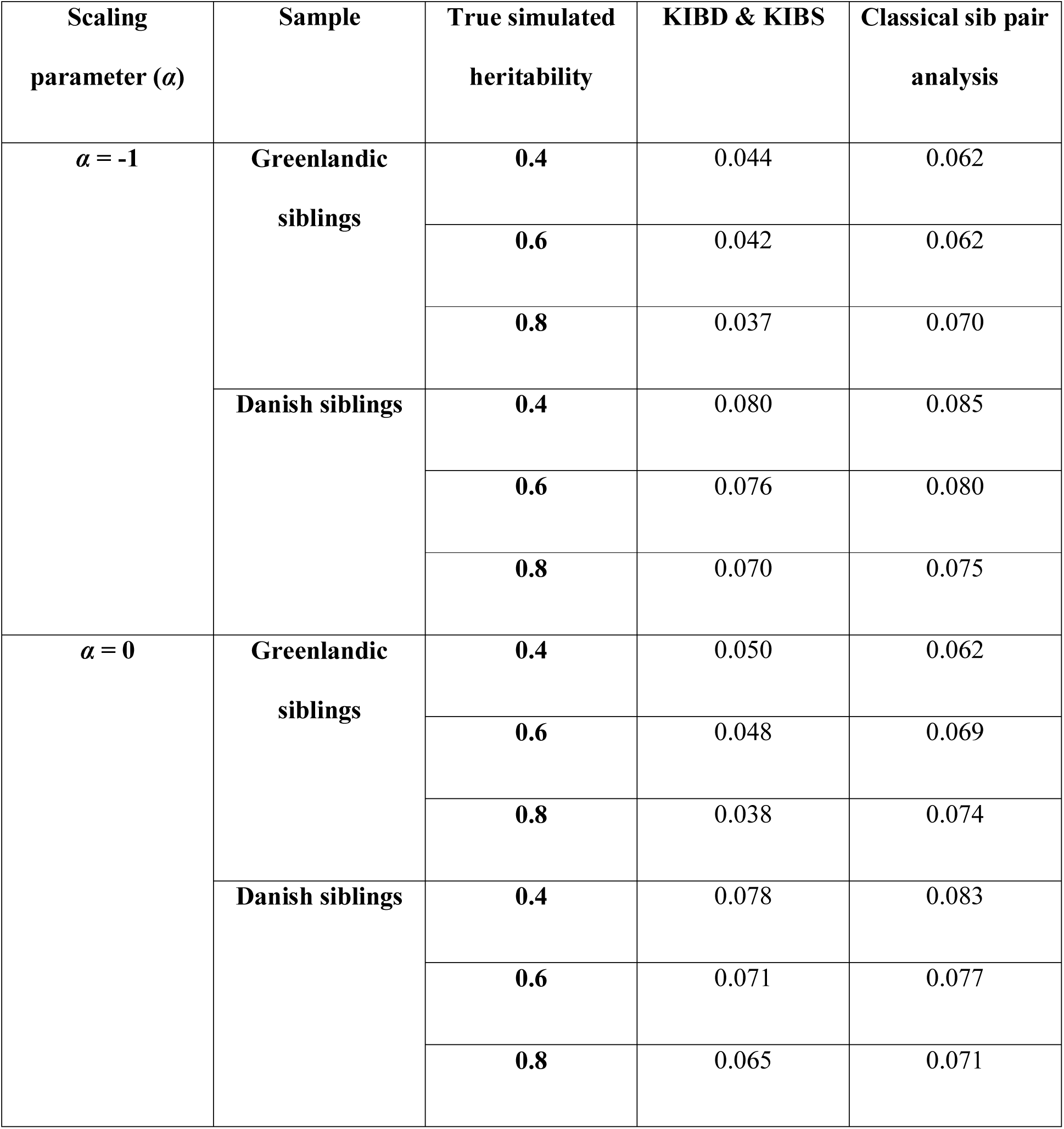
root-mean-square deviation of the total heritability estimates from their respective true simulated values for the **K**_**IBD**_ & **K**_**IBS**_ and the classical sib-pair analysis

**Table S2:**
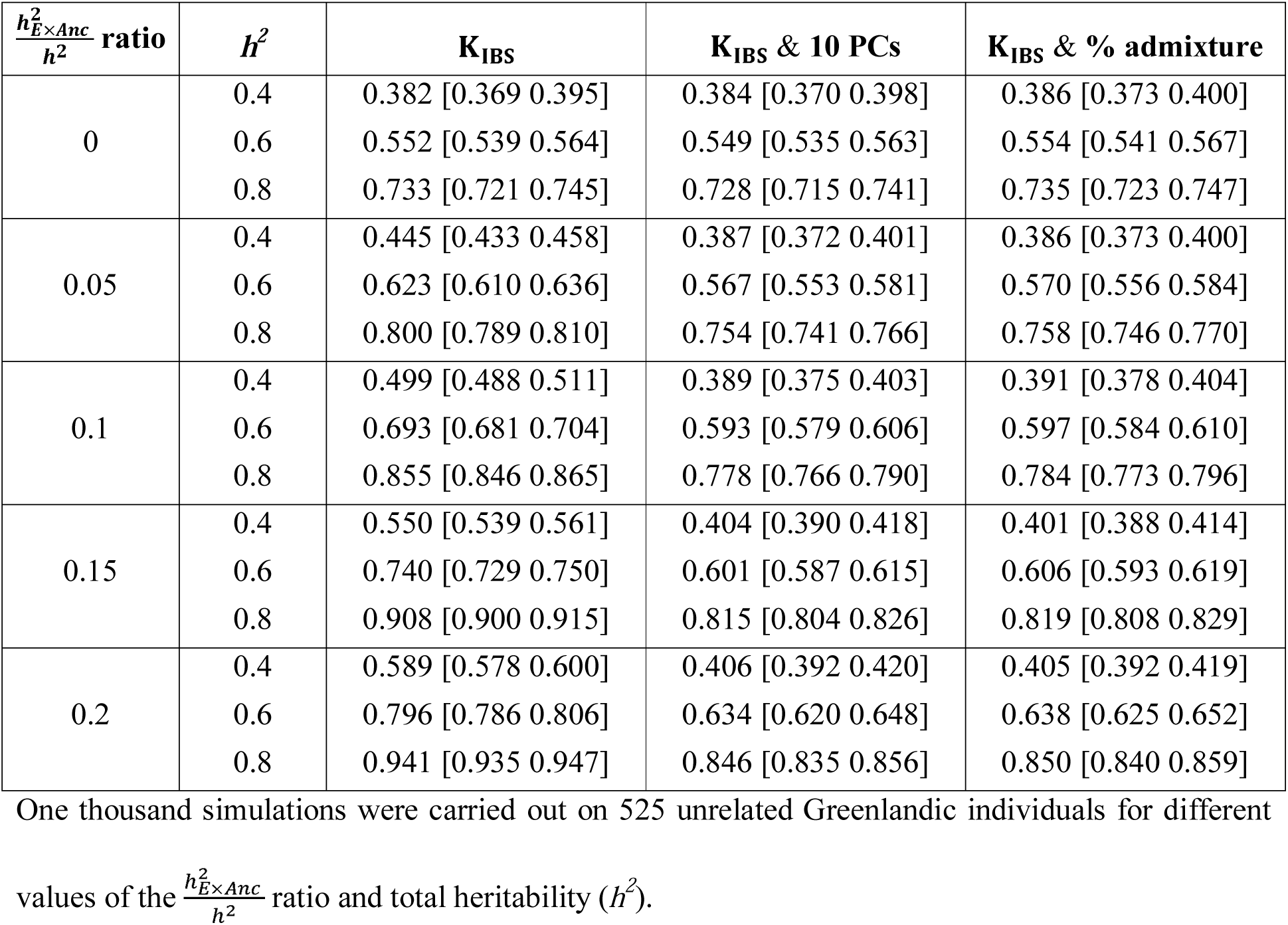
mean SNP heritability 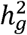 estimates and 95% confidence intervals (in square brackets) under the **K**_**IBS**_ model without and with covariates

**Table S3:** total heritability estimates of the ten quantitative traits from the Greenlandic sib pairs using the **K**_**IBD**_ & **K**_**IBS**_ & age & sex & 10 PCs model under the LDAK model assuming different values for the genotype scaling parameter *α*

| Alpha ($\alpha$) | Height | Weight | BMI | Hip | Waist | Waist-to-hip ratio | Cholesterol | HDL cholesterol | LDL cholesterol | Triglycerides |
| --- | --- | --- | --- | --- | --- | --- | --- | --- | --- | --- |
| -1.25 | 0.766±0.041 | 0.501±0.049 | 0.438±0.049 | 0.522±0.049 | 0.463±0.050 | 0.377±0.053 | 0.512±0.049 | 0.469±0.049 | 0.524±0.052 | 0.317±0.053 |
| -1 | 0.781±0.041 | 0.513±0.049 | 0.446±0.050 | 0.527±0.049 | 0.469±0.051 | 0.382±0.053 | 0.520±0.050 | 0.485±0.049 | 0.531±0.052 | 0.322±0.053 |
| -0.75 | 0.788±0.041 | 0.523±0.049 | 0.454±0.050 | 0.531±0.050 | 0.474±0.051 | 0.384±0.054 | 0.523±0.050 | 0.498±0.049 | 0.535±0.052 | 0.326±0.053 |
| -0.5 | 0.789±0.041 | 0.530±0.049 | 0.460±0.050 | 0.533±0.050 | 0.477±0.051 | **0.385±0.054** | 0.524±0.050 | 0.506±0.050 | 0.536±0.052 | 0.329±0.054 |
| -0.25 | **0.786±0.041** | 0.533±0.049 | 0.465±0.050 | 0.533±0.050 | 0.478±0.051 | 0.383±0.054 | **0.523±0.050** | 0.510±0.050 | 0.535±0.052 | 0.331±0.054 |
| 0 | 0.781±0.041 | 0.535±0.049 | 0.468±0.050 | 0.534±0.050 | 0.479±0.051 | 0.382±0.053 | 0.521±0.050 | 0.511±0.049 | **0.534±0.052** | 0.333±0.054 |
| 0.25 | 0.776±0.041 | **0.536±0.049** | **0.470±0.050** | **0.534±0.050** | **0.479±0.051** | 0.380±0.053 | 0.519±0.050 | **0.512±0.049** | 0.533±0.052 | **0.333±0.054** |

Values in bold indicate the estimate with the highest likelihood.

